# The Arousal-Regulated Filter: Modulating Feedforward and Recurrent Dynamics for Adaptive Neural Tracking

**DOI:** 10.1101/2025.03.31.646280

**Authors:** Jonathon R. Howlett

## Abstract

Classic behavioral studies have identified an inverted-U relationship between arousal and performance, while neurophysiology studies have shown that the arousal-related neuromodulator norepinephrine (NE) increases the signal-to-noise ratio (SNR) of neural responses to extrinsic inputs. More recently, abstract computational models suggest that arousal signals uncertainty in predictive internal models and increases the influence of new observations. Here, I present the arousal-regulated filter (ARF), a novel computational model of neural state estimation designed to bridge abstract algorithms, behavioral findings, and basic neural mechanisms. According to the ARF, arousal selectively amplifies feedforward synapses relative to recurrent synapses, consistent with findings from neurophysiology. Computationally, the ARF integrates predictions of an internal model, implemented in recurrent connections, with extrinsic observations, relayed by feedforward projections, to adaptively track dynamic systems. The ARF resembles a Kalman filter but replaces dynamic updating of the Kalman gain matrix with modulation of a scalar gain parameter influenced by arousal, and incorporates a potentially nonlinear activation function. Computational simulations demonstrate ARF’s versatility across binary, multi-unit categorical, and continuous neural network architectures that track diffusion and drift-diffusion processes. When the internal model is aligned with environmental dynamics, arousal exhibits an inverted-U relationship with accuracy due to a bias-variance tradeoff, consistent with behavioral results. However, when the internal model is misaligned with the true dynamics (i.e. the true state is unpredictable), increased arousal monotonically improves accuracy. Optimal arousal levels vary systematically, being lower in noisy sensory contexts and higher in volatile environments or when unmodeled dynamics exist. Thus, the ARF provides a unifying framework that links abstract computational algorithms, behavioral phenomena, and neurophysiological mechanisms. The model also offers potential insights into the computational effects of altered arousal states in mental health conditions, with potential implications for new approaches to assessment and treatment.

**Author Summary:** To effectively navigate the world, the brain must integrate internal predictions with new sensory information. One factor believed to affect this balance is arousal, which controls alertness and attentiveness to new information. Models of the effect of arousal on the brain have tended to be either very abstract (concerning abstract mathematical algorithms) or very concrete (concerning the effect of arousal-related neurotransmitters like norepinephrine on electrical activity in neurons), but the link between these two levels of analysis has not been fully clear. In this study, I introduce the arousal-regulated filter (ARF), a new model designed to bridge abstract computation and concrete neural mechanisms. The model proposes that arousal modulates the influence of new information by altering the balance of different types of neural connections. Through a series of computer simulations of different neural networks, I show that moderate arousal provides a balance between accuracy and responsiveness, which may be disrupted by overly low or high arousal. Overall, this work may help integrate previous theories of arousal and help understand disruptions of arousal in mental illness such as anxiety and posttraumatic stress disorder (PTSD).

## Introduction

In order to accurately represent the world, organisms must combine noisy sensory observations with predictions from internal models. Recent theoretical and empirical work has demonstrated that this process is regulated by the arousal response, mediated by neurotransmitters such as norepinephrine (NE) [1, 2]. NE and arousal are thought to signal a greater level of uncertainty in internal model predictions, and therefore to increase emphasis on new observations relative to predictions [1].

Adaptive fluctuations in arousal are therefore critical in maintaining efficient neural computation in the face of shifting environmental context and task demands.

In parallel to the development abstract theoretical models of the role of arousal in neural computation, empirical investigations have detailed the effects of NE on cellular and synaptic functions. However, the findings of this body of research have generally resisted simple, overarching summaries. NE has diverse effects on synaptic function, either increasing or decreasing the gain on excitatory synapses depending on neural regions and cortical layers [3]. It is therefore an important challenge to reconcile the diverse, nonuniform synaptic and cellular effects of NE with the putative overarching role of the arousal response in neural computation.

A classic result regarding the effect of NE on neural function is the finding that NE increases signal-to-noise (SNR) ratio of neuronal responses to extrinsic inputs [4]. An explanation of this effect was offered by Hasselmo et al., who found that NE significantly suppresses excitatory recurrent synapses in the rat piriform cortex, while having a weaker effect in suppressing excitatory afferent synapses [5]. This suggests that the “noise” which is suppressed by NE may in fact represent a computationally meaningful contribution of recurrent connections to neural activity. Moreover, a relative effect of NE in suppressing recurrent synapses compared to feedforward synapses could be an important facet of its overall computational function.

This paper presents the arousal-regulated filter (ARF), a novel model linking the synaptic effects of NE (and potentially other arousal-related neurotransmitters) to a core neurocomputational function. Specifically, the model posits that neural state estimation combines a predictive model implemented in recurrent connections with new observations relayed via feedforward connections. Similar to the Kalman filter and other state estimation algorithms [6], the predictive model implemented in recurrent connections allows the ARF to “filter” noisy observations. Similar to the Kalman gain, arousal increases the gain on feedforward inputs relative to the gain on recurrent connections, thus achieving more optimal state estimation when observations have high reliability relative to the internal model. Using ARF simulations, I illustrate the role of arousal in reducing bias at the expense of increasing variance in neural tracking of dynamic systems. The results suggest that ARF may unify findings from the electrophysiology literature, in which high levels of NE improve neural SNR, with behavioral findings encapsulated in the Yerkes-Dodson law, in which there is an inverted-U relationship between arousal and performance. The key insight is that increasing arousal increases accuracy in state estimation in the setting of a misaligned internal model (e.g. for an unpredictable extrinsic input in an electrophysiology experiment), but that arousal yields an inverted-U relationship with accuracy in the setting of an aligned internal model (due to the bias-variance tradeoff). Consistent with behavioral findings, the optimal arousal level depends on the nature of the tracking task, being lower with high input noise and higher with high process noise (i.e. volatility) or unmodeled dynamics. The results support the ARF as a model which links abstract computational and algorithmic functions to concrete neural implementation, potentially fostering a more detailed understanding of the role of arousal in health and disease.

## Results

### The Arousal-Regulated Filter (ARF)

The ARF performs state estimation by combining new observations with the predictions of an internal state transition model. The state which is estimated by the ARF may represent a multivariate latent state, enabling abstraction from multivariate afferent inputs. The state is represented as a vector, which may be implemented as activations of a set of neural units in the state layer. Separately, the afferent input is represented by an observation vector which may be implemented as activations of afferent input neurons. The feedforward synapses are represented as a feedforward weight matrix which maps the observation space onto the state space (which may have different dimensions).

The internal state transition model is implemented by recurrent synaptic connections within the state representation layer. In the absence of new observations, the state vector evolves as a dynamical system, which represents the predicted evolution of the true environmental state. The set of recurrent synaptic weights is represented as a recurrent weight matrix. In the brain, the predictive model can include projections from other brain regions in addition to local recurrent connections within a region. Formally, these distant projections and synapses can be included in the state vector and recurrent weights within the ARF. Arousal modulates state estimation in the ARF by increasing the weight on feedforward connections relative to recurrent connections. This increases the influence of new observations relative to model predictions.

The ARF equation is:

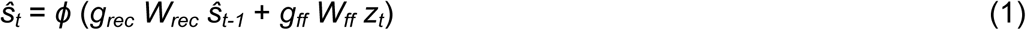

where *ŝ*_t_ is the state vector estimate at time *t*, *ϕ* is a (potentially nonlinear) activation function inspired by neural activation functions, *g*_rec_ is a scalar gain on recurrent synapses, *W_rec_* is the matrix of recurrent connection weights, *ŝ_t-1_* is the state vector estimate at time *t-1, g_ff_* is a scalar gain on feedforward synapses, *W_ff_* is the matrix of feedforward connection weights, and *z_t_* is the new observation vector.

### Relationship Between the ARF and the Kalman Filter

The ARF exhibits two important differences from the Kalman filter: 1) dynamic updating of the Kalman gain matrix based on process noise covariance and measurement covariance is replaced by dynamic updating of the two scalar gain terms *g*_rec_ and *g_ff_*; 2) the potentially nonlinear activation function *ϕ* enables nonlinearities in the prediction and observation models, unlike the Kalman filter (although it should be noted that this does not guarantee optimal nonlinear state estimation).

The ARF can be shown to be equivalent to a simplified Kalman filter with a fixed Kalman gain matrix, under the conditions that *ϕ* is the identity function and the weight matrices *W_rec_* and *W_ff_* are chosen appropriately. We start with the Kalman prediction step equation:

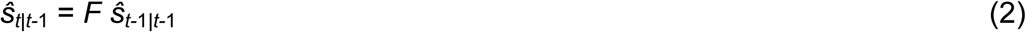

where *ŝ_t|t-_*_1_ is *ŝ_t_* (the state vector estimate at time *t*) predicted using information up to time *t-*1, *ŝ_t-_*_1*|t-*1_ is *ŝ_t-_*_1_ (the state vector estimate at time *t-*1) predicted using information up to *t-*1, and *F* is the state transition matrix, and the Kalman state update equation:

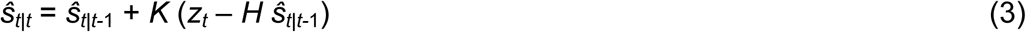

where *K* is the Kalman gain and *H* is the observation matrix. Substituting (2) into (3), we have:

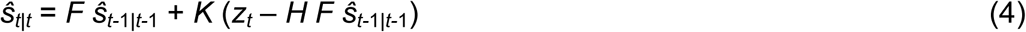

Rearranging:

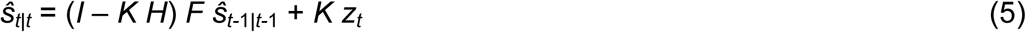

which is equivalent to the ARF in which *ϕ* is the identity function, *g_rec_ W_rec_*= (*I* – *K H*) *F*, and *g_ff_ W_ff_* = *K*.

Alternatively, the ARF can be applied to a predictive coding framework in which feedback projections carry predictions from the network to the input layer, and the feedforward projections from the input layer represent prediction errors rather than observations. In this case, we can substitute a prediction error term (*z_t_* – *H F ŝ_t-_*_1_) for the observation term *z_t_* in our ARF equation:

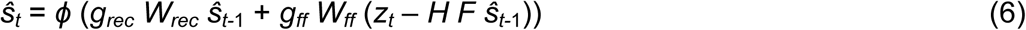

where *H*, the observation matrix, now represents a feedback weight matrix translating the prediction *F ŝ_t-_*_1*|t-*1_ from the state space to the observation space where it is subtracted from the observation *z_t_*. This is equivalent to equation (4) if *ϕ* is the identity function, *g_rec_ W_rec_* = *F*, and *g_ff_ W_ff_* = *K*.

### Two-Unit Binary Network Simulations

The ARF was applied to simulate a simple network which could take one of two discrete values, labeled “0” and “1” (Fig. 1). The network was comprised of two binary units representing the two possible states as network activities (1, 0) and (0, 1). At each timestep, the network combined noisy inputs (via feedforward connections) with a prediction based on the previous state (via recurrent connections) to determine the next state. The feedforward weight matrix was set as the identity matrix so that unit 0 would be stimulated when the true state was 0 and unit 1 would be stimulated when the true state was 1. Independent Gaussian noise was added to each input unit (i.e. the input unit for the correct state was computed as 1 plus Gaussian noise and the input unit for the incorrect state was computed as 0 plus Gaussian noise). The recurrent connections included positive self-connections and negative cross-connections:

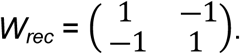

**Figure 1:**
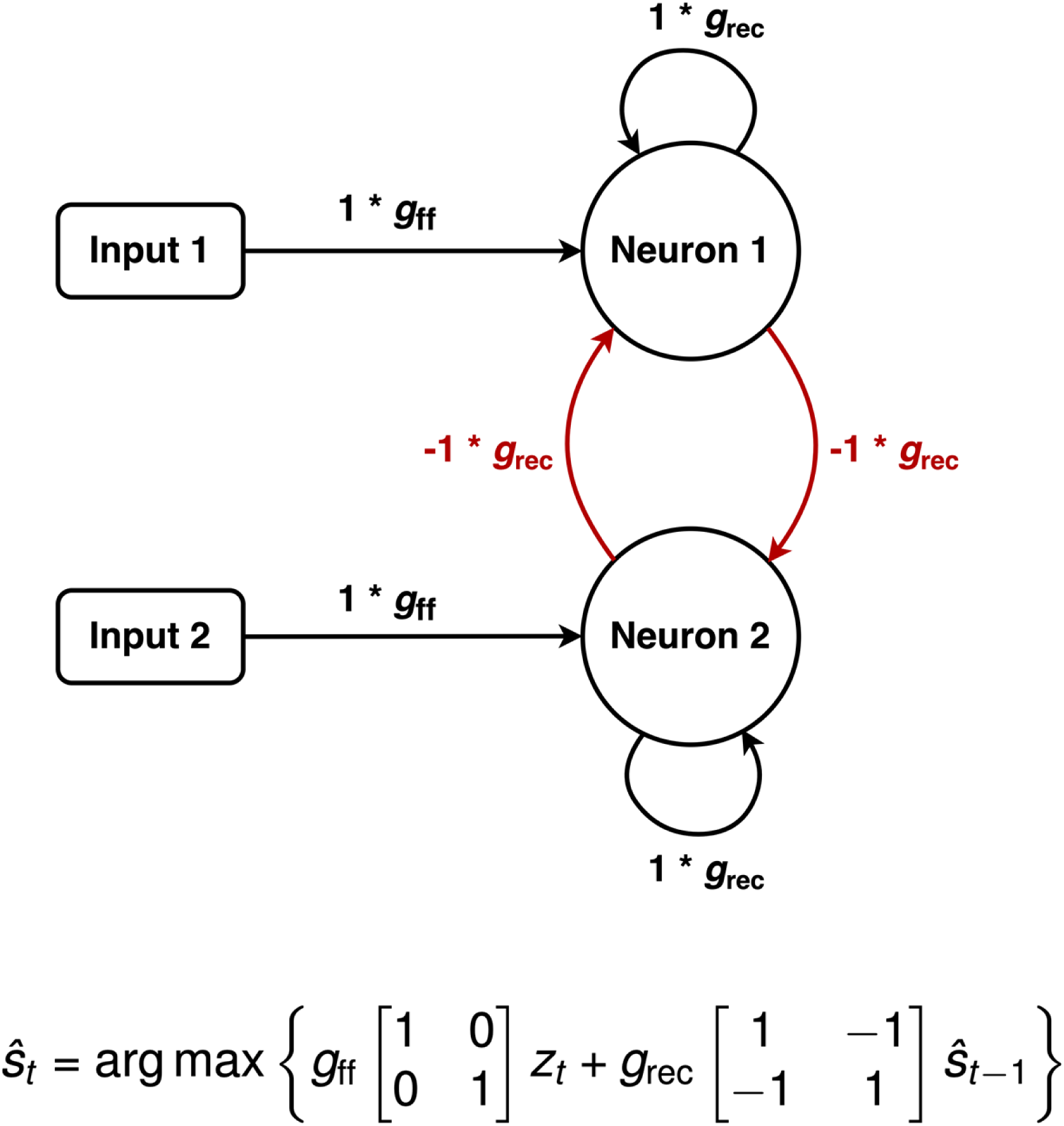
Arousal-Regulated Filter Applied to Binary Neural Network. A two-neuron network which can take one of two discrete values, with each neuron representing one state. The network combines noisy inputs (via feedforward connections) with a prediction based on the previous state (via recurrent connections) to determine the next state. The feedforward weight matrix for this network is the identity matrix. Recurrent connections include positive self-connections and negative cross- connections, so the internal model predicts the state will remain stable for the next time step. Arousal controls the balance between feedforward weights (scaled by *g_ff_*) and recurrent weights scaled by *g_rec_*). An argmax operation enforces a binary state space.

Arousal was operationalized as a scaling factor multiplied by the feedforward weight matrix. An argmax operation was performed at each timestep (i.e., the unit with the higher net input was set to 1 and the other unit was set to 0), in order to enforce a binary state space.

First, simulations were performed in which the network was initialized to state 0 and the true state was set at state 1, to determine the effect of arousal on the network’s response to a true change in state (see Fig. 2a). As expected, networks with higher arousal responded to a change in state more quickly. Networks with higher arousal also plateaued in mean network state value earlier than networks with lower arousal.

**Figure 2:**
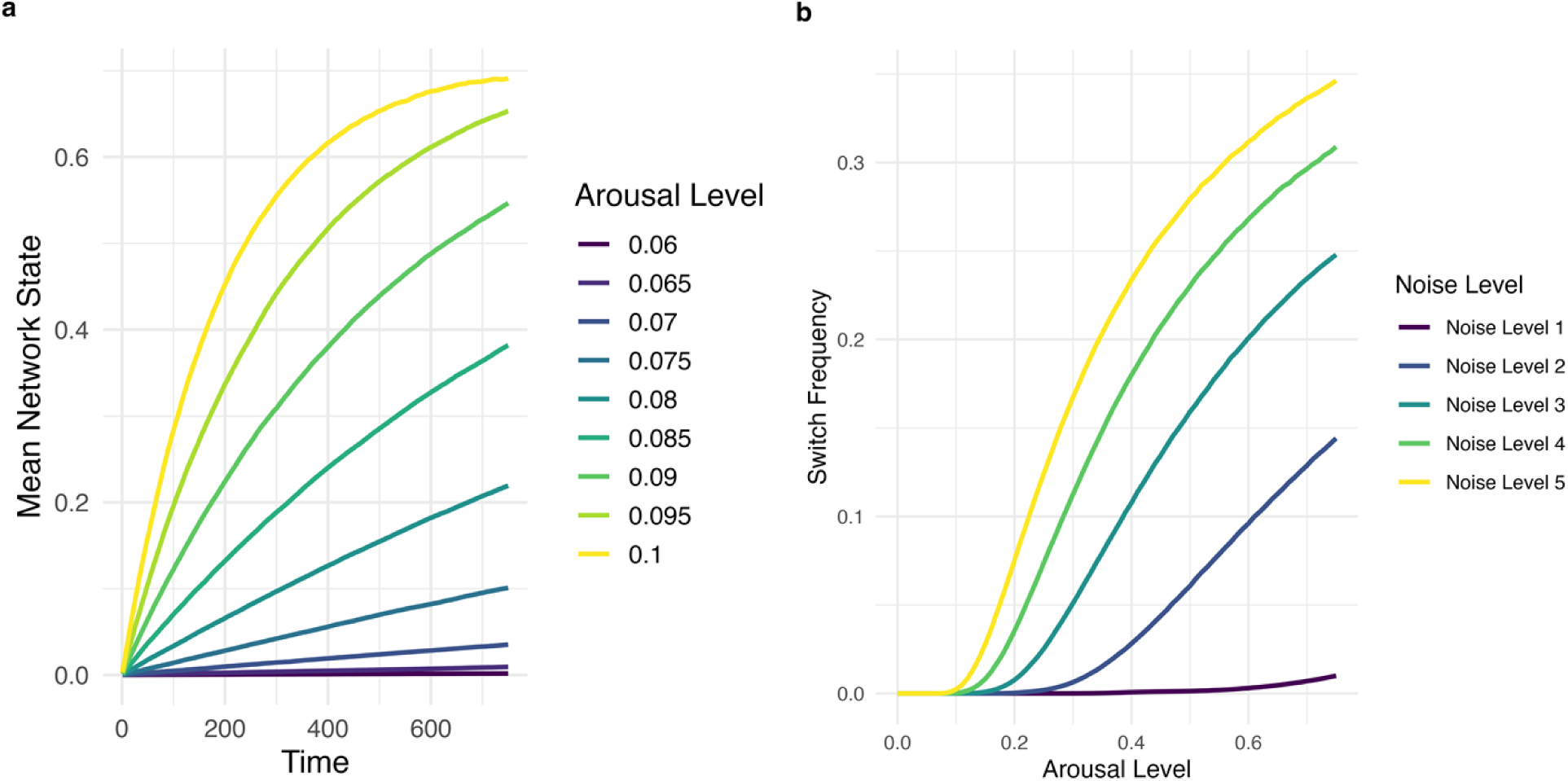
Arousal Increases Network Responsiveness and Decreases Network Stability. (a) Network response after a true change in state. The binary network is initialized to state 0 and the true state is set to state 1. Mean network state is plotted over time for varying levels of arousal. Higher arousal levels cause a faster network response after the change in state. (b) Effect of arousal and measurement noise on switch frequency. Frequency of network state change is plotted for varying arousal and measurement noise for an unchanging true state (hazard rate 0). Higher arousal and higher measurement noise lead to reduced stability of the network state.

Next, simulations were performed in which the true state changed at intervals based on a hazard rate, i.e. the probability of state change at each timepoint. Arousal, noise, and hazard rate were systematically varied to examine their effects on both the switch frequency of the network (i.e. how often the network changed state; see Fig. 2b) as well as the accuracy of the network (i.e. the proportion of the time the network was in the correct state; see Fig. 3). As expected, switch frequency increased with both increasing noise level and increasing arousal (as higher arousal amplified the feedforward noise). Switch frequency was plotted for hazard rate 0 only because switch frequency was similar across varying arousal levels. As expected, there was an inverted-U relationship between accuracy and arousal. At an arousal level of 0, in which activity was solely driven by recurrent connections, accuracy was at chance level (with the proportion of time the network was in the correct binary state being 0.5). As arousal increased and therefore the influence of feedforward input increased, accuracy began to exceed chance level, rose to a peak, and then declined. This finding reflects a balance in the tradeoff between two effects of higher arousal: faster response to a true change and higher switch rate in the absence of a true change (meaning that state representations are less stable with higher arousal). The shape of the inverted-U relationship between arousal and accuracy was influenced by both hazard rate and noise, such that higher arousal levels were optimal for higher hazard rates and lower noise levels. With higher hazard rates, the faster response rate with high arousal becomes more important, while with higher noise, the higher switch rate with higher arousal becomes more important, again reflecting the varying tradeoff between rapid tracking and stability.

**Figure 3:**
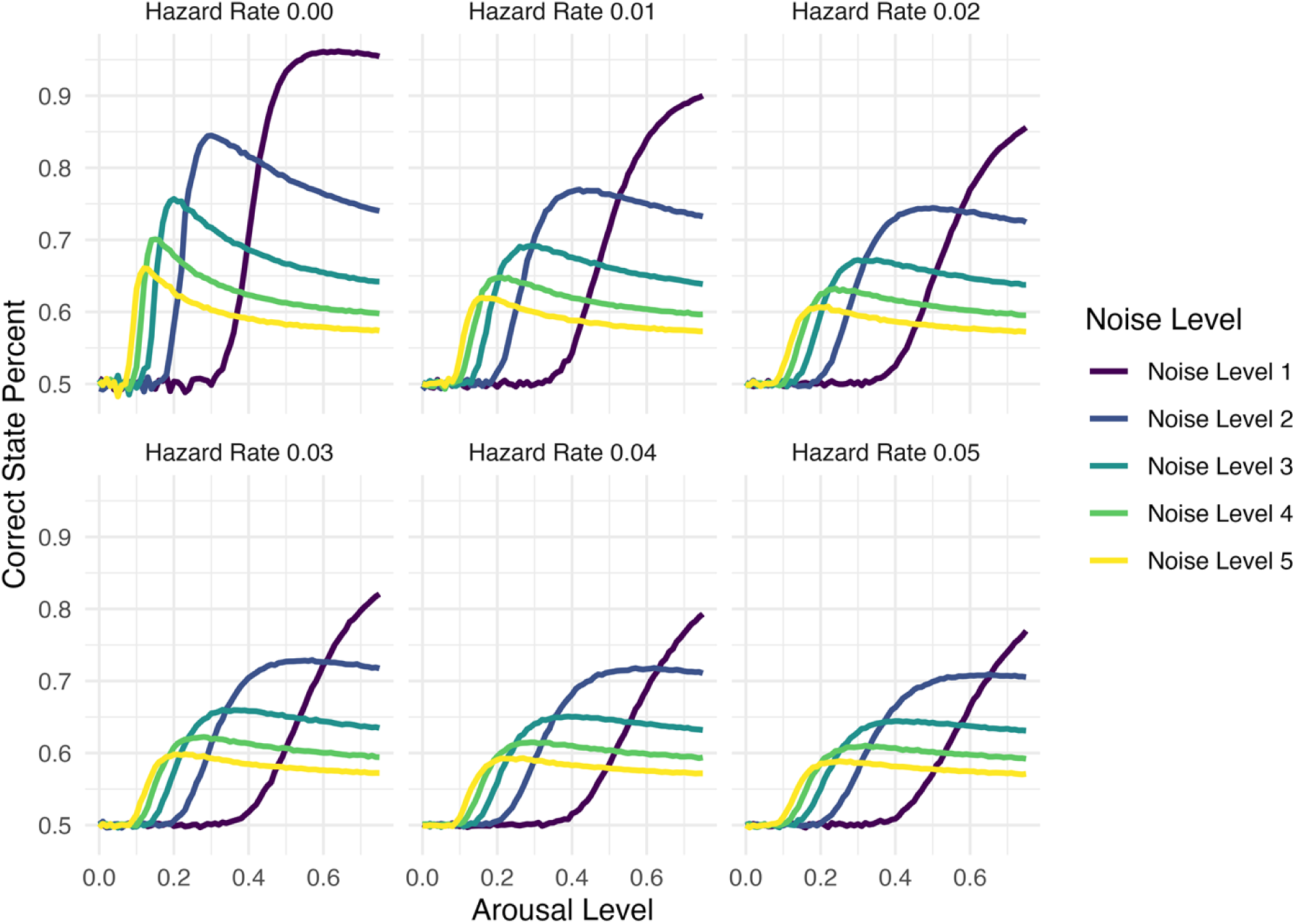
Effect of Arousal, Measurement Noise, and Hazard Rate on Accuracy. Mean accuracy (proportion of time in the correct state) plotted for varying levels of arousal, measurement noise, and hazard rate (rate of change of the true state) for the binary network. Arousal displays an inverted-U relationship with accuracy. Peak accuracy occurs at higher arousal levels for lower measurement noise and higher hazard rate.

### Multi-Unit Categorical Network

In the preceding simulations, the recurrent connections favored staying (i.e. remaining in the previous state) over switching. Thus, the internal dynamic model of the network was consistent with the true state dynamics when the true hazard rate was less than 0.5 (although the network did not explicitly represent the probability of switching).

Additionally, the argmax operation enforced a binary state space, meaning the network was not allowed to represent “impossible” states such as (0, 0) and (1, 1), which were not part of the true state space. In order to explore network performance when the internal network model was misaligned with the true dynamics, a more complex network was constructed which allowed impossible states and enabled construction of recurrent weight matrices that were either congruent or incongruent with the true system dynamics.

The network was constructed with 12 binary units. The feedforward matrix was a 12-dimensional identity matrix, i.e. each network unit received a single feedforward connection from a particular input unit. The recurrent weight matrix exhibited excitatory connections within the first six units and within the last six units, and other connections being inhibitory. This tended to favor the states (0, 0, 0, 0, 0, 0, 1, 1, 1, 1, 1, 1) and (1, 1, 1, 1, 1, 1, 0, 0, 0, 0, 0, 0), although other states were allowed. The true state space was binary and was either congruent with the internal network model, with states (0, 0, 0, 0, 0, 0, 1, 1, 1, 1, 1, 1) and (1, 1, 1, 1, 1, 1, 0, 0, 0, 0, 0, 0), or incongruent with the internal network model, with states (0, 0, 0, 1, 1, 1, 0, 0, 0, 1, 1, 1) and (1, 1, 1, 0, 0, 0, 1, 1, 1, 0, 0, 0). See Fig. 4a for a diagram depicting the congruent and incongruent networks.

**Figure 4:**
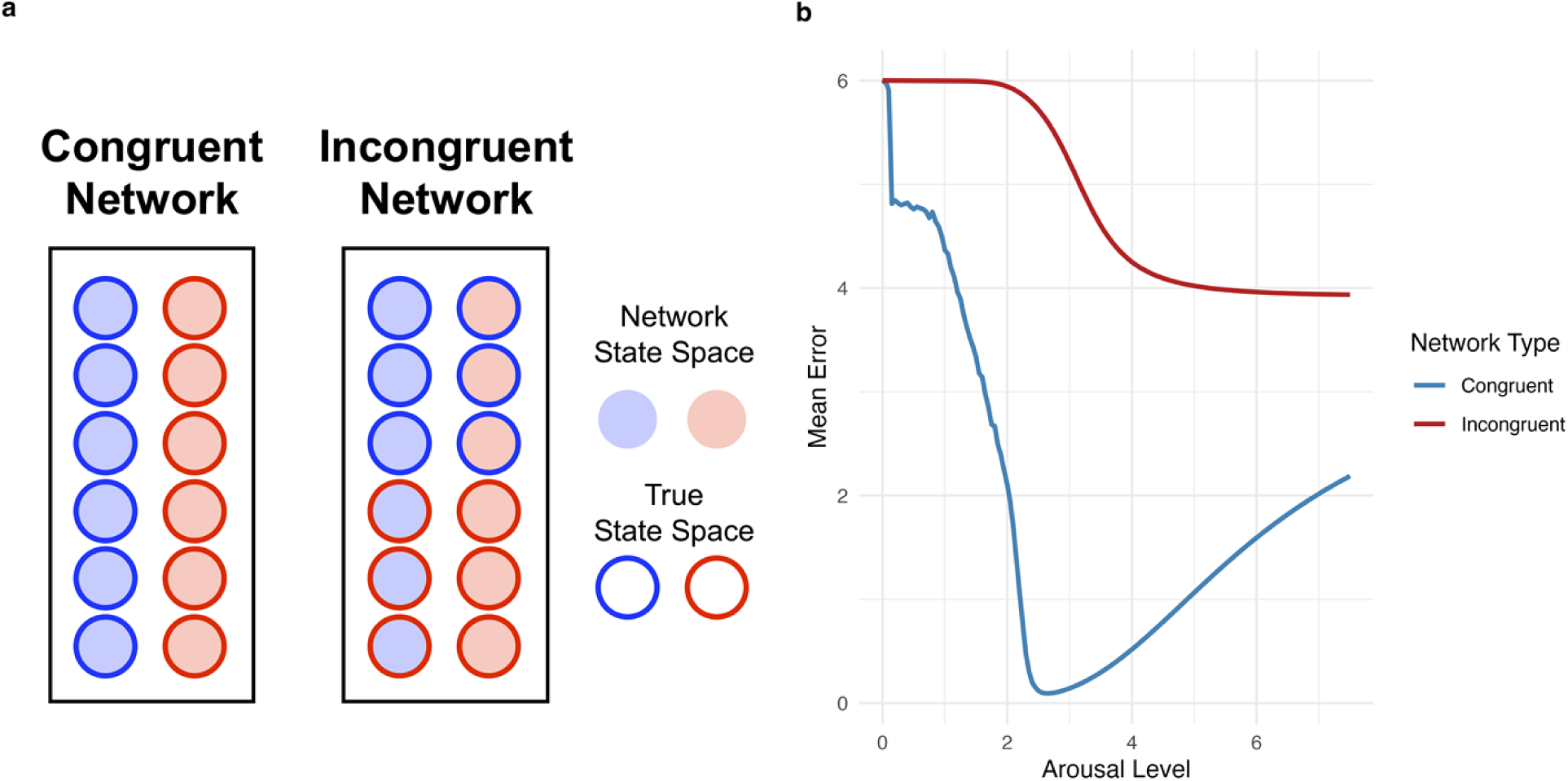
Effect of Arousal Depends on Alignment of Internal Model with Environment. (a) Congruent and incongruent multi-unit categorical networks. Twelve-unit networks encode a preferred binary state space via their recurrent connections (excitatory between units shaded the same color and inhibitory between units shaded different colors). The true binary state space, with states relayed via noisy feedforward inputs to each neuron, is either congruent or incongruent with the preferred network state space. (b) Effect of arousal in congruent and incongruent networks. Mean error is plotted for various arousal levels for congruent and incongruent networks. For the congruent network, arousal displays a U-shaped relationship with error due to the tradeoff between responsiveness to the true state relayed by feedforward connections and the stabilizing effect of recurrent connections. The sudden drop with a slight increase in arousal above 0 is due to initial states which are halfway between the two attractor states. For the incongruent network, error monotonically decreases with increasing arousal because the misaligned internal model is not useful in stabilizing state representations.

The true state was randomly selected from the two possible states and was continued throughout each run (i.e. hazard rate was zero). Gaussian noise was added separately to each of the 12 input units (i.e. Gaussian noise was added to 0 or 1 for each input unit depending on the true state). Arousal was operationalized as a scaling factor multiplied by the feedforward identity matrix. Noise (i.e. standard deviation of the Gaussian noise added to the input units) was fixed at 1. Both the congruent and the incongruent network were simulated with varying levels of arousal with randomized starting states, and the error (i.e. number of mismatches between the true state vector and the network vector) was measured at each timepoint (see Fig. 4b).

At arousal 0, when activity was determined entirely by recurrent connections, both networks performed at chance level (i.e. mean error of 6 out of 12 units). For the congruent network, there was a steep reduction in error with a slight increase in arousal above 0, which reflected starting states halfway between the attractors for the two possible states being pushed toward the correct state with a small amount of feedforward input. Then, there was continued reduction in error with increasing arousal until a minimum was reached, followed by a subsequent increase in error. This U-shaped curve reflects the tradeoff between the need for feedforward input to indicate the true state and the benefit of the congruent recurrent weights for filtering noise. In contrast, the incongruent network evidenced a reverse-S relationship between arousal and error, indicating that increasing arousal continued to reduce error indefinitely because the incongruent recurrent weight matrix was not useful in improving state estimates. At very high levels of arousal, the curves for the congruent and incongruent networks asymptotically approach each other as the influence of the different recurrent weights continues to decline.

### Continuous Network Tracking of Random Diffusion Dynamics

A neural network was simulated with a single continuous unit that tracked a time-varying true state. The true state evolved with diffusion dynamics, i.e. at each timepoint random Gaussian process noise (volatility) was added to the current position. A feedforward projection relayed the true position to the position unit in the network, with Gaussian measurement noise added. The neural state evolved according to the equation:

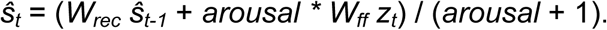

Runs were conducted with varying levels of process noise (volatility), measurement noise, and arousal (Fig. 5a). Both the network and the true process were initialized to position 0 at the beginning of each run. Except for the case of a volatility of 0 (in which case the optimal arousal level was 0), there was a U-shaped relationship between arousal and mean absolute error in position. Minimum error occurred at higher levels of arousal for higher volatility and at lower levels of arousal for higher noise.

**Figure 5:**
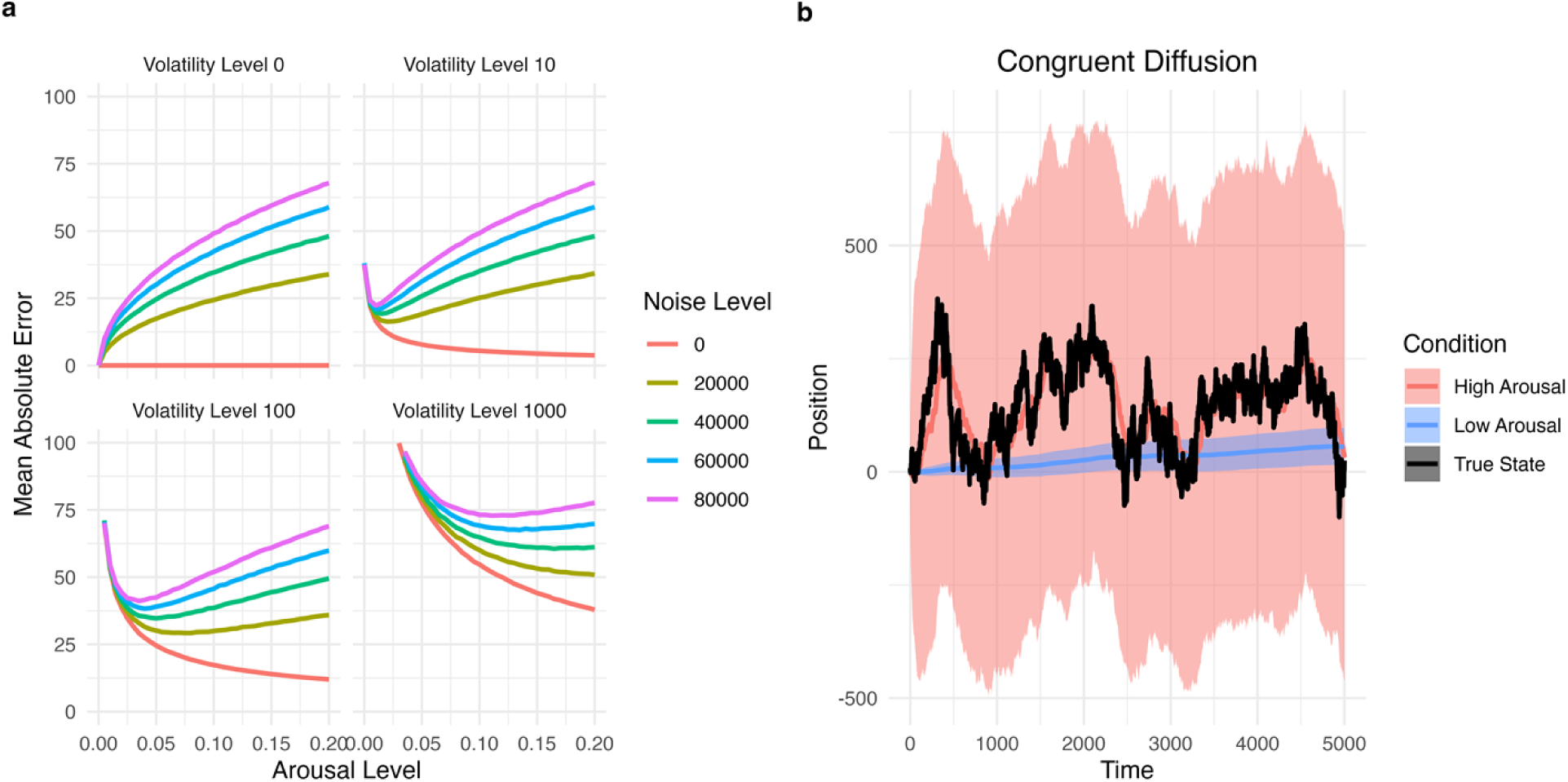
Continuous Network Tracking of Diffusion Dynamics. (a) Mean absolute error plotted for varying levels of arousal, measurement noise, and volatility (or process noise) for a continuous network tracking a state with diffusion (i.e. random walk or Brownian motion) dynamics. For volatility level 0, error increases with increasing arousal (because the network was initialized in the correct state). For positive volatility, arousal displays a U-shaped relationship with error. Higher arousal is optimal for lower levels of measurement noise and for higher levels of volatility. (b) Bias-variance tradeoff for tracking diffusion dynamics. A single trajectory of true states was generated, and then separate input sequences were generated by adding independently sampled measurement noise. Mean (colored line) and standard deviation (shaded ribbon) were then plotted across runs for high and low arousal levels. The high arousal level displays a low level of bias (mean network state closely tracks true state) but a high level of variance (the standard deviation across runs is large). The low arousal level displays a higher level of bias but a low level of variance. The tradeoff between bias and variance determines the optimal arousal level for the network.

Next, the bias-variance tradeoff was examined for the diffusion tracking problem (Fig. 5b). Diffusion networks with two different arousal levels (low and high) were applied to a true diffusion process with fixed volatility and measurement noise. First, a single true state trajectory was generated which was used for all subsequent runs.

Next, for each run, an input sequence was generated by adding newly sampled Gaussian measurement noise to the true state trajectory. This input sequence was then fed to one of the networks. For each network, mean estimated state was plotted, along with the standard deviation of estimated state across runs. The purpose of this procedure was to distinguish bias, the systematic difference between the true state trajectory and the mean network estimate, from variance, the additional random error contributed by measurement noise. As expected, the low-arousal network exhibited high bias but low variance while the high-arousal network exhibited low bias but high variance. This occurred because increased arousal reduces the stabilizing influence of recurrent connections, causing the network to rely more heavily on noisy feedforward inputs. Thus, while high arousal corrects systematic bias, it also increases variance by amplifying measurement noise.

### Network Tracking of Drift-Diffusion Dynamics

In order to apply the ARF to a more complex dynamic system, simulations were performed in which continuous networks tracked a state generated by drift-diffusion dynamics. At each timepoint a drift rate (velocity) term was added to the current position, along with Gaussian process noise (or volatility). For these simulations, drift rate was fixed, i.e. process noise applied to position but not to drift rate. The neural networks consisted of 2 continuous units, the first of which tracked position and the second of which tracked drift rate. The recurrent weight matrix implemented drift-diffusion dynamics in the internal model:

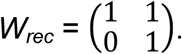

A feedforward projection relayed the true position to the position unit in the network, with Gaussian measurement noise added. The neural state evolved according to the same equation as in the diffusion simulations.

Two networks were simulated: an incongruent network in which the network drift rate was set to 0 and a congruent network in which the network drift rate was set to the true drift rate for the generating process. This was done to examine the effect of unmodeled dynamics, i.e., a drift term in the generating process for the true state that was not included in the network’s internal model. These two networks were simulated for varying arousal levels (Fig. 6a). Both networks displayed a U-shaped relationship between arousal and mean absolute error. The congruent network had a lower minimum error, and minimum error occurred at a lower arousal level. This indicates that greater arousal is needed to overcome unmodeled dynamics in the incongruent case by amplifying feedforward inputs.

**Figure 6:**
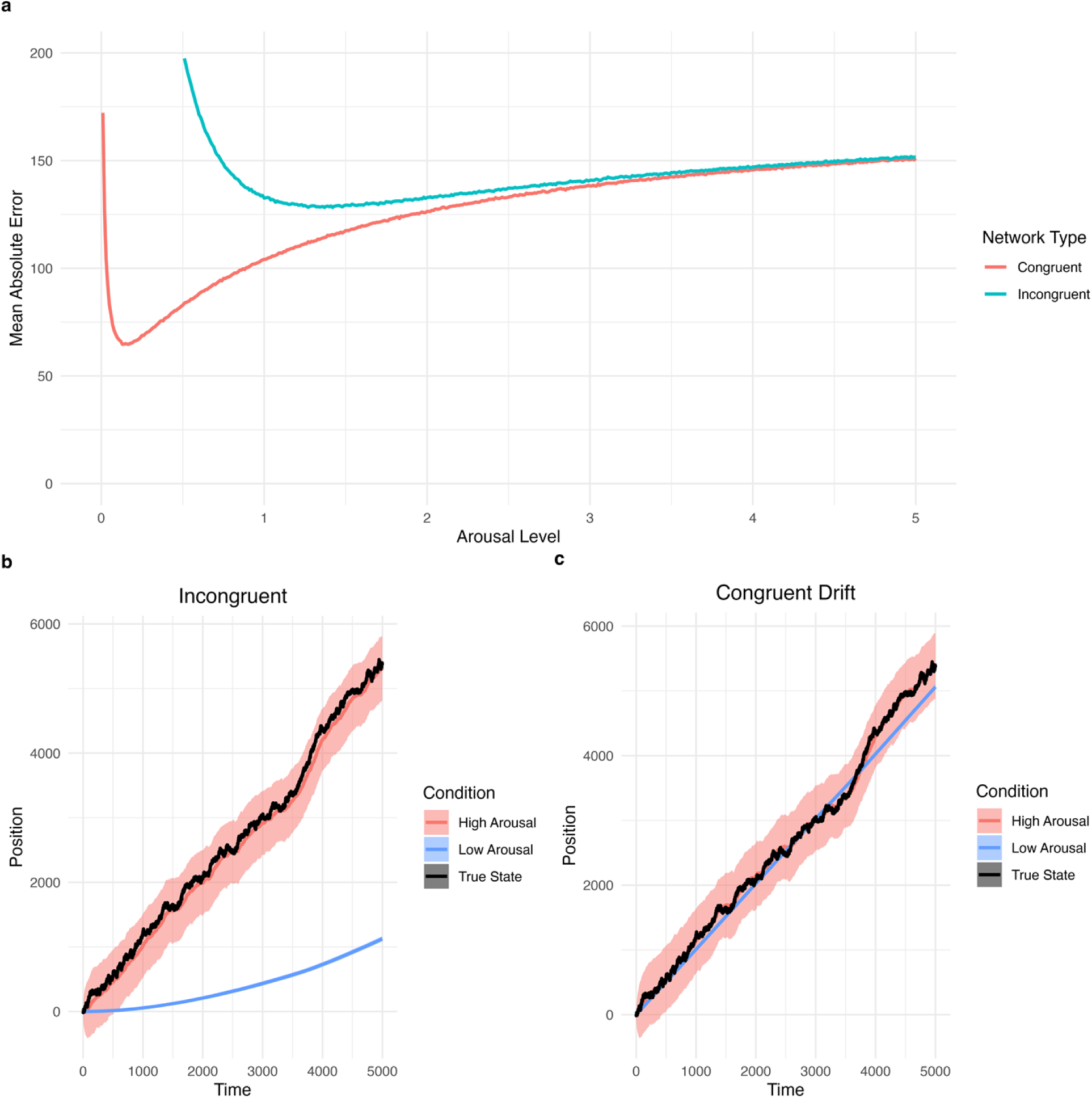
Continuous Network Tracking of Drift-Diffusion Dynamics. (a) Effect of arousal in congruent and incongruent continuous networks tracking drift-diffusion dynamics (i.e. biased random walk). The incongruent network does not incorporate a drift rate term into its internal model (i.e. recurrent connections only model random diffusion dynamics). The congruent network incorporates drift-diffusion dynamics with the internal drift rate set to the true drift rate of the generating process. Optimal arousal is higher for the incongruent network due to the need to overcome unmodeled dynamics via stronger feedforward inputs. (b) Bias-variance tradeoff for incongruent network. With low arousal, the incongruent network displays a high level of bias as it does not anticipate the positive drift of the true generating process. High arousal is needed to overcome the unmodeled dynamics to reduce bias, at the expense of higher variance due to amplified measurement noise. (c) Bias-variance tradeoff for congruent network. For the congruent network, relatively low bias is achieved with a low arousal level because the network’s internal model anticipates the positive drift of the generating process. Thus, by incorporating the true dynamics into the predictive model, both low bias and low variance are achieved (resulting in overall lower mean error).

Next, bias-variance analyses were performed for both the congruent and incongruent networks. As before, a single true state trajectory was generated for use in all runs, while separate input sequences were generated by adding newly sampled measurement noise for each run. Each network was simulated for both low arousal and high arousal conditions. In the incongruent case, in which the internal network model reflected diffusion dynamics while the true state evolved according to drift-diffusion dynamics, low arousal caused a high degree of bias as the network lagged far behind the true state (Fig. 6b). High arousal led to low bias, as the mean network state closely tracked the true state, at the expense of high variance due to amplification of measurement noise. In contrast, in the congruent case, in which the network evolved according to drift-diffusion dynamics that matched the generating process for the true state, low arousal resulted in relatively low bias while retaining low variance (Fig. 6c). Thus, a better-performing internal predictive model allowed the network to track the state evolution with reduced need for feedforward input, reducing the effect of measurement noise in destabilizing the state representation.

### Adaptive Drift Rate Tracking (Predictive Coding Implementation of an Alpha-Beta Filter)

While the preceding simulations involved a fixed network drift rate, the ARF is also capable of adaptively tracking the drift rate based only on position input (i.e., when the true drift rate is not directly observed). To illustrate this, a network was constructed which adaptively estimates drift rate using an alpha-beta filter [7] implemented in a predictive coding framework (Fig. 7a). In this network, rather than reflecting an observation, the feedforward projections relay prediction errors. The current predicted position is fed back to the input layer, where it is subtracted from the position observation. This position prediction error is then projected both to the position unit and the drift rate unit in the network, with the feedforward weights being scaled by arousal. The projection weight to the position neuron is equivalent to the alpha parameter in an alpha-beta filter, while the projection weight to the drift rate neuron is equivalent to the beta parameter. As before, the recurrent weight matrix reflected a drift-diffusion process:

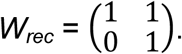

**Figure 7:**
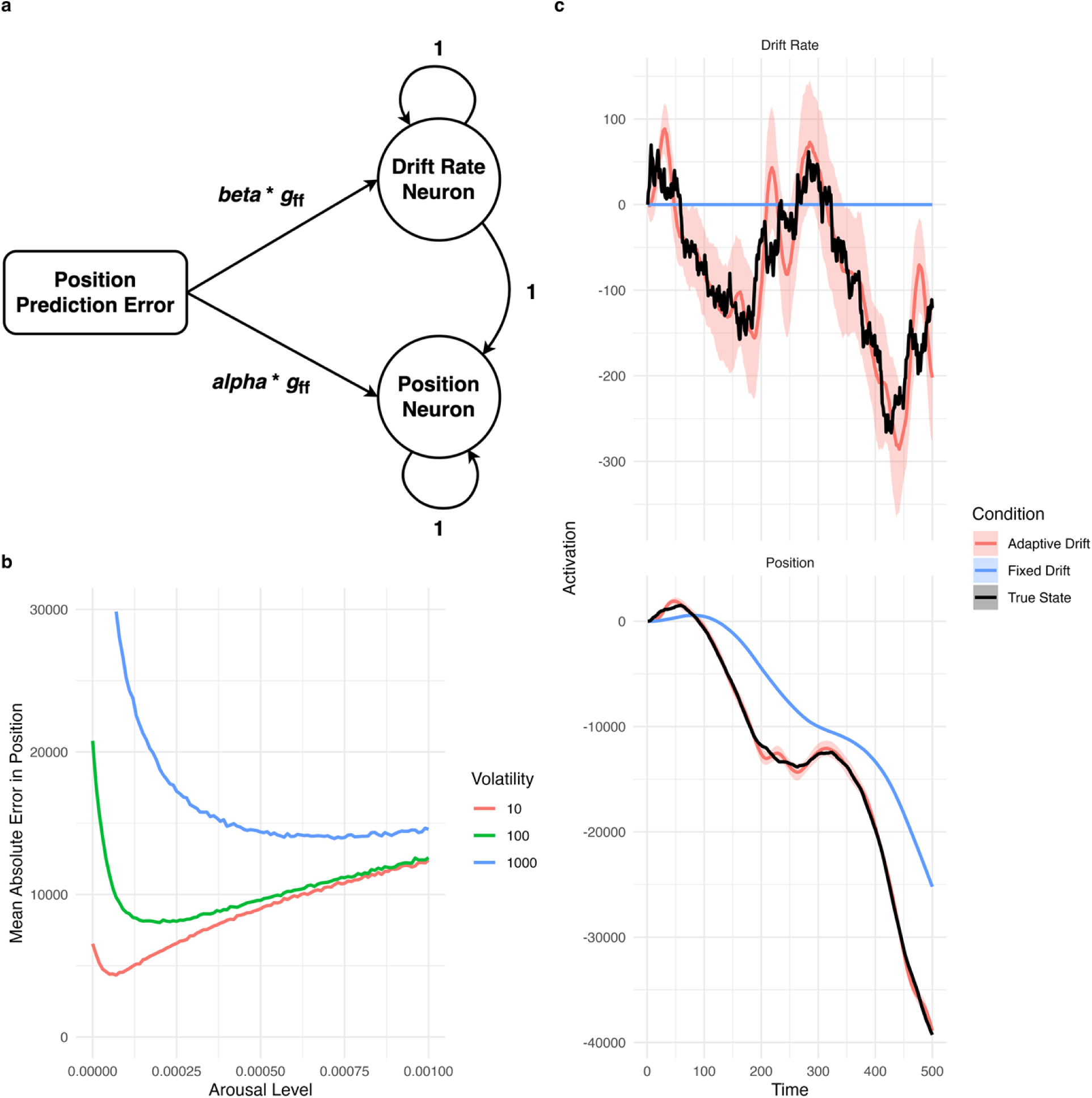
Alpha-Beta Filter Implemented in Predictive Coding Network. (a) Network implementation of alpha-beta filter for adaptive tracking of drift rate. Rather than having a fixed drift rate, the adaptive drift network continually estimates the true drift rate based only on observed position at each time point. Prediction error in position is fed to both the position unit and the drift rate unit, scaled by fixed alpha and betaarameters and by arousal level. Feedback connection relaying predicted position from the network to the input layer is not shown. Recurrent weights implement the drift-diffusion prediction model. (b) Effect of arousal and volatility in adaptive drift network. Arousal displays a U-shaped relationship with error. Optimal arousal is achieved at a higher level for higher levels of volatility (or process noise). Process noise affects both the position and the drift rate (i.e. the drift rate varies randomly over time). (c) Adaptive drift and fixed drift networks. The adaptive drift network tracks the varying true drift rate over time based only on prediction error in position (i.e. the true drift rate is not observed directly by the network). As a result, the adaptive drift network displays lower bias in tracking the true position compared to a fixed drift network with the same arousal level.

As before, the true state was generated by a drift-diffusion process. However, in these simulations, process noise was applied to both position and drift rate, so that the drift rate of the true process varied over time. The state for both the network and the true process was initialized to (0, 0).

To examine the role of arousal in tuning the alpha and beta parameters of the alpha-beta filter, the adaptive drift network was simulated under varying levels of arousal and volatility (process noise). There was a U-shaped relationship between arousal and mean absolute error, with higher arousal levels being optimal for higher volatility (Fig. 7b). This suggests that the alpha and beta parameters of the alpha-beta filter can be tuned by arousal to minimize tracking error based on the rate of unpredictable change in the underlying system. In the ARF framework, the alpha and beta parameters are both linearly scaled by arousal rather than being differentially tuned by arousal, but the ratio between the alpha and beta parameters could be altered through synaptic learning.

Simulations were performed for two networks: an adaptive drift network, as described above, and a network with drift rate fixed at 0 (fixed drift network). Bias-variance analyses were performed, with a single true state trajectory being generated and used across all runs, and with separate input sequences generated for each run by adding newly sampled Gaussian noise. Results revealed that the adaptive drift network tracked the state much more closely than the fixed drift network for an equivalent level of arousal (Fig. 7c).

## Discussion

The ARF is a model of neural state estimation, and its modulation by arousal-related neuromodulators such as NE, which is designed to bridge abstract computational models and algorithms [1], behavioral findings (such as those encapsulated in the Yerkes-Dodson law [8]), and the electrophysiology literature (which suggests that NE differentially alters the gain on feedforward and recurrent synapses [5]). The ARF proposes that arousal selectively amplifies feedforward inputs, which convey noisy observations or prediction errors, relative to recurrent connections, which implement an internal, predictive dynamic model. The flexible nature of the ARF enables application to a variety of simple network types and dynamic systems, including binary networks, multi-unit categorical networks, and continuous tracking networks.

These simulations reveal several major features of the influence of arousal on state estimation. First, arousal increases the speed of response to unpredicted changes in state while reducing the stability of internal state representations. Second, arousal reduces bias in state tracking by amplifying the influence of new observations, at the expense of increasing variance by amplifying measurement noise. Third, arousal exhibits a U-shaped relationship with error (or inverted-U relationship with accuracy) when the internal predictive model is aligned or partially aligned with the true system dynamics, but exhibits a monotonically decreasing relationship with error when the internal model is misaligned. Finally, optimal arousal is lower with higher measurement noise and is higher with higher process noise or volatility (i.e., unpredictable true system change) and when unmodeled dynamics are present. Taken together, these findings indicate that the ARF may serve as an integrative framework for understanding the role of arousal in state estimation across a range of cognitive processes and for linking this computational role to synaptic mechanisms.

ARF is positioned as a bridge model between abstract computational mechanisms, behavioral and cognitive findings, and cellular and synaptic research on the effects of neuromodulators such as NE. This bridging perspective may help guide future empirical studies based on theoretical predictions. For example, ARF provides a theoretical framework for interpreting the diverse effects of NE across cortical regions and layers (by positing that a critical feature of NE action may be its effect on feedforward synapses relative to recurrent synapses rather than its effect on feedforward synapses per se). Thus, while the isolated observation that NE suppresses the gain on feedforward inputs to the piriform cortex may obscure its computational role, the finding that it suppresses the gain on recurrent synapses to a greater degree may be critical in clarifying this role. A strong theoretical model could help guide investigation of the role of NE in other neural regions, potentially resolving diverse findings into a coherent framework and grounding the effects of arousal across cognitive processes in specific neural implementations.

Beyond guiding neurophysiology research, ARF may also help resolve seemingly contradictory findings across behavioral and neural data. Classical behavioral results point to an inverted-U relationship between arousal and performance, while classical electrophysiology results indicate that increased NE improves the neural SNR for an extrinsic cortical input. The ARF may resolve this contradiction by distinguishing between aligned and misaligned internal models. When the internal model is misaligned with the true dynamics, as in the incongruent categorical network simulations, error in state representations decreases monotonically with increasing arousal because recurrent connections no longer play a useful filtering role. This may be the case in electrophysiology research in which a sudden, unpredictable extrinsic input is experimentally generated. In these studies, the contribution of recurrent synapses in modulating the response to feedforward inputs would contribute to “noise” in the neural SNR. However, in behavioral contexts in which an animal is performing a cognitive task, these same recurrent synapses may be aligned with the dynamics of the system being tracked play an important role in filtering measurement noise. In this case, the optimal balance between feedforward and recurrent connections will depend on the bias-variance tradeoff as influenced by measurement noise, volatility or process noise, and unmodeled dynamics. As suggested in the categorical network simulations, misalignment between the internal network model may take the form of an incongruence between the network’s preferred state space and the true environmental state space. The network’s state space may represent a neural manifold, or a low-dimensional set of patterns within a higher dimensional network which represent a state space enforced by recurrent connections [9]. The current results suggest that high levels of arousal may “pull” network representations off this low-dimensional manifold to represent input features which are incongruent with the internal network state space, and that arousal may therefore regulate the dimensionality of neural representations by modulating the influence of feedforward and recurrent connections.

The ARF may offer additional insight into the effects of arousal state on behavior.

Accounts of the of locus coeruleus (LC), which distributes NE throughout the central nervous system, emphasize the distinction between states of low tonic LC activity, in which animals are drowsy and poorly responsive to stimuli, moderate tonic LC activity, in which animals respond quickly to task-relevant stimuli, and high tonic LC activity, in which animals are prone to distractibility and rapid changes in behavior [10]. These effects may have substantial clinical relevance for disorders characterized by excessive arousal or distractibility, such as anxiety disorders [11], posttraumatic stress disorder (PTSD) [12], and attention deficit hyperactivity disorder (ADHD) [13]. The ARF may help to further explicate the computational and neural mechanisms of the influence of arousal on behavior in healthy populations and in mental health disorders. Future application of the ARF may help to better assess and treat alterations in arousal and their specific influence on cognitive and behavioral performance.

The ARF is related to optimal state estimation algorithms such as the Kalman filter, but also differs in several key respects. In the Kalman filter, the Kalman gain is a matrix which maps prediction errors in the observation space into the state space. This matrix is adaptively updated at each timestep based on the covariance matrices for measurement noise and for uncertainty in the current state prediction [6]. In contrast, the ARF involves a fixed feedforward weight matrix which is adaptively scaled by a scalar gain term which depends on arousal. This constraint is inspired by the low-dimensional nature of LC signals which drive release of NE throughout the brain but is likely an oversimplification as NE release may be locally modulated [14]. The scalar gain term implies that the ARF is suboptimal for many filtering problems. A second key distinction between the Kalman filter and the ARF is the potentially nonlinear activation function, inspired by neural activation functions [15], which enables application of the ARF to nonlinear dynamic models. However, it is important to note that this does not guarantee optimality in nonlinear state estimation. Future work can further integrate the ARF with more detailed models of nonlinear neural filtering, including accounts of how neural filters are learned through synaptic mechanisms [16].

Future work can apply the ARF to specific cognitive processes, rather than the simple, abstract dynamic systems simulated in this manuscript. The original Yerkes-Dodson findings concerned a visual discrimination task with different discrimination difficulties [8], which could be readily operationalized by varying the signal-to-noise ratio of extrinsic inputs into a categorical network, as in the simulations performed here.

However, the role of arousal in complex processes such as motor control, working memory, and cognitive control could be modeled with further network extensions. For example, the cerebellum is thought to implement a dynamic forward model in motor control tasks [17], and the ARF can be easily extended by positing that cerebellar projections, along with local recurrent connections, are attenuated by arousal relative to feedforward inputs. In addition, neural gating mechanisms in working memory [18] and cognitive control [19] tasks may partially overcome the tradeoff between speed of updating and the stability of representations by controlling the timing of extrinsic inputs to prefrontal circuits.

Limitations of the ARF include the lack of biological realism of its simple network models (which is intentional given the role of ARF as a bridging model between abstract computational algorithms and biological implementation). Future work will be necessary to explore ARF-like computation in more biologically plausible network models, including spiking neurons with membrane time constants, interneuron populations, transmission delays, and other features of biological networks [15]. The ARF also does not explicitly consider synaptic learning or the role of arousal in learning [20] and is not intended to represent a full theory of the effects of arousal on neural processing. The ARF is also focused on the downstream effects of arousal rather than its upstream drivers (such as the regulation of arousal by the anterior cingulate (ACC) and orbitofrontal cortices (OFC) [10]), although it may enrich understanding of these processes by further clarifying the optimal level of arousal under a particular set of computational circumstances.

In conclusion, the ARF represents a promising approach to integrate abstract computational models and algorithms such as the Kalman filter, studies of the effect of arousal on cognition and behavior, and neurophysiologic studies of the effect of neuromodulators such as NE on neural processing. The ARF can be flexibly applied to state estimation in a range of dynamic systems and could be extended to incorporate it into more detailed models of cognitive processes or biologically detailed networks. The model may be useful in better understanding the computational role of arousal and its neural implementation and could be applied the assessment and management of disorders involving dysregulation of arousal.

## Methods

### Two-Unit Binary Network

All simulations were performed in R [21]. For the simulations in which the two-state network responded to a true change in state, each run was 750 timepoints long and conducted with randomized Gaussian noise with standard deviation of 5 added to input units. Runs were conducted with varying arousal levels (operationalized as a scaling term multiplied by the feedforward identity matrix), from 0.06 through 0.1, with increments of 0.005. A total of 100,000 runs were conducted for each arousal level, and the mean values across runs were plotted.

For the simulations in which the true state changed based on a hazard rate, simulation runs were 500 timepoints long. Noise levels (i.e. the standard deviation of the Gaussian noise added to input units) were varied from 1 to 5, incremented by 1. Hazard rates were varied from 0 to 0.05, varied by 0.01. A total of 5,000 runs were conducted for each parameter combination, with mean values across runs being plotted.

### Multi-Unit Categorical Network

The recurrent weight matrix for the multi-unit categorical network was:

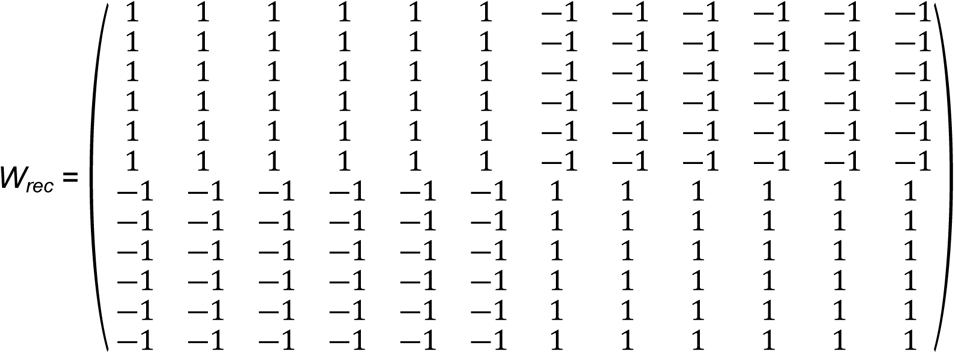

The networks were simulated for arousal levels from 0 to 7.5, in increments of 0.05, with each run consisting of 500 timesteps. Each network was simulated for 50,000 runs at each arousal level. Mean error was calculated for each network type at each arousal level.

### Continuous Network Tracking of Diffusion and Drift-Diffusion Dynamics

For the simulations in which drift rate was set to 0 for both the network and the true generating process and mean absolute error was plotted, runs were simulated with measurement noise levels from 0 to 8,000, incremented by 2,000 and with volatility levels of 0, 10, 100, and 1000. Runs were simulated for arousal levels from 0 to 0.2, incremented by 0.005. Each run was 500 timepoints long. For each combination of parameters, 2,000 runs were conducted, and mean absolute error in the position estimate was calculated across runs.

For the bias-variance analysis of the diffusion process, the low arousal value was set to 0.0001 and the high arousal value was set to 0.01. Volatility (i.e. process noise) was set to 100. Measurement noise was set to 50,000,000. Simulations were run for 5,000 timepoints. For each combination of parameters, 1,000 runs were conducted.

For the congruent and incongruent network comparison for the drift-diffusion process, the true drift rate was set at 100. Simulations were run at arousal levels from 0 to 5, in increments of 0.01. Volatility (i.e. process noise) was set at 1,000 and measurement noise was set at set at 50,000. Simulations were run for 500 timepoints. For each combination of parameters, 500 runs were conducted.

For the bias-variance analysis of the drift-diffusion process, the low arousal value was set to 0.0001 and the high arousal value was set to 0.01. True drift value was set to 1. Volatility (i.e. process noise) was set to 100. Measurement noise was set to 50,000,000. Simulations were run for 5,000 timepoints. For each combination of parameters, 1,000 runs were conducted.

### Predictive Coding Implementation of an Alpha-Beta Filter

For the simulations comparing the adaptive drift network to the fixed drift network, process noise (volatility) was set to 100 for both position and drift rate. Measurement noise was set to 1,000. Arousal was set at 0.01. For the adaptive drift network, the feedforward matrix (before being scaled by arousal) was:

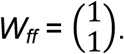

For the fixed drift network, the feedforward matrix was:

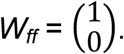

For each network, 50 runs were conducted.

For the simulations of the adaptive drift network with varying arousal and process noise (volatility), runs were performed for arousal levels from 0 to 0.002, incremented by 0.00001 and for volatility levels of 10, 100, and 1000. Measurement noise was set to 50,000. Each run consisted of 500 timepoints. For each combination of parameters, 5,000 runs were conducted.

## Acknowledgements

This research was supported by the Veterans Health Administration Office of Research and Development Career Development Award IK2 CX001887 (to JRH).

